# SimuCell3D: 3D Simulation of Tissue Mechanics with Cell Polarization

**DOI:** 10.1101/2023.03.28.534574

**Authors:** Steve Runser, Roman Vetter, Dagmar Iber

**Affiliations:** Department of Biosystems Science and Engineering (D-BSSE), ETH Zürich, Schanzenstrasse 44, 4056 Basel, Switzerland; Swiss Institute of Bioinformatics (SIB), Schanzenstrasse 44, 4056 Basel, Switzerland

## Abstract

The 3D organisation of cells determines tissue function and integrity, and changes dramatically in development and disease. Cell-based simulations have long been used to define the underlying mechanical principles. However, large computational costs have so far limited simulations to either simplified cell geometries or small tissue patches. Here, we present SimuCell3D, a highly efficient open-source program to simulate large tissues in 3D with subcellular resolution, growth, proliferation, extracellular matrix, fluid cavities, nuclei, and non-uniform mechanical properties, as found in polarised epithelia. Spheroids, vesicles, sheets, tubes, and other tissue geometries can readily be imported from microscopy images and simulated to infer biomechanical parameters. Doing so, we show that 3D cell shapes in layered and pseudostratified epithelia are largely governed by a competition between surface tension and intercellular adhesion. SimuCell3D enables the large-scale *in silico* study of 3D tissue organization in development and disease at an unprecedented level of detail.

## Introduction

The acquisition and maintenance of proper morphology are crucial for the normal physiological functioning of a biological tissue. Their disruptions are associated with a range of pathological conditions, including cancer and birth defects. The shape of tissues is determined by the dynamic positioning of their constituent cells, which can collectively deform or migrate to induce macroscopic changes in the tissue morphologies [1, 2]. These cellular behaviors are regulated by the mechanical properties of both the cells and extracellular matrix (ECM) [3], along with the distribution of stresses within tissues [4]. Therefore, understanding how tissues acquire and maintain their shapes requires a deep comprehension of the interplay between the stress distribution within them and the mechanical properties of their cells and ECM.

Various experimental methods have been developed to contribute to this understanding [5–7], *e*.*g*., micropipette aspiration [8], atomic force microscopy [9], optical stretcher [10], and laser ablation [11]. Nonetheless, these experimental techniques are generally limited to the rare tissues directly accessible to probing, or to small tissue portions. In addition, even when all the factors influencing a tissue morphology have been experimentally identified, their synergy might remain unclear.

Recent advances in the fields of fluorescent microscopy, image processing, and computation power now allow to complement these direct measurements with *in silico* models, and thus to gain a more global understanding of the cellular dynamics underlying tissue morphogenesis and homeostasis [12–17]. Among these numerical methods, cell-based models have become widely used in the fields of developmental and cancer biology due to their high spatio-temporal resolution and accurate predictions. Cell-based models recreate virtual versions of tissues by representing cells as individual agents with their own mechanical properties and behavior. These models offer an *in silico* environment where the stress distribution and the mechanical properties of cells can be modulated to study their impact on tissue morphology and function. Cell-based models can thus predict the tissue shape arising from experimentally measured cell properties or, conversely, in conjunction with parameter estimation methods, they can allow to infer the cell properties that led to an imaged tissue morphology. The high level of spatio-temporal details of cell-based models, however, entails a substantial computational cost which forces them to a tradeoff between the number of cells they can simulate and the spatial resolution of their representation [18]. For this reason, cell-based models with varying levels of resolution have been developed to address different types of biological problems. For instance, center-based models are a class of cell-based models that represent cells as simple spheres, making them suitable for phenomena where the abundance of cells is more crucial than their shape. These models have been used to gain deeper understanding of a wide range of phenomena including, for instance, the development of tumors [19] or the inflammation of tissues [20].

Vertex models are another class of cell-based model that have been developed to study tissues in which cell shapes can be approximated by polygons in 2D [21–25] or polyhedrons in 3D [26]. This simplification allows them to represent each cell with only a few points, enabling them to simulate large tissues. Vertex models have been employed to study a wide range of phenomena, including the transition between solid-like and fluid-like tissue states [27], as well as various morphogenetic processes such as the polarization of early embryos [28], the formation of branched structures [29], and the biased elongation of tissues [30, 31]. However, their simplistic representation of tissues comes with the drawback that they cannot adequately represent cells with complex shapes. Furthermore, the highly restricted topology permissible for the mesh in vertex models significantly complicates the simulation of phenomena like cell extrusion or tissue fusion. The mechanisms underlying these developmental events are among the fundamental open problems in morphogenesis.

To address the limitations of vertex models, a family of cell-based models sometimes referred to as Deformable Cell Models (DCMs) has been developed. These models provide a more geometrically realistic representation of tissues by discretizing each cell membrane separately into a closed loop of connected points in 2D [40–51] or a closed triangulated surface in 3D [32–39, 52, 53]. The complex shapes cells can adopt in DCMs make these models particularly suited to simulate phenomena such as the development of early embryos [54] or the cellular movements during wound healing [35]. However, the accuracy offered by these models comes at a staggering computational cost. To mitigate this computational cost, one 3D DCM implementation named CellSim3D [36] constrains the cell geometries to spheroidal shapes. This approach is however not suited for the study of tissues with complex (non-polyhedral) cell shapes. The remaining 3D DCMs preserve a high geometrical resolution of the cell membranes but are limited by their computational efficiency. At best, they can simulate the growth of a tissue from one to a thousand cells in a week of computation time [35], precluding their use for large-scale computational studies. Additionally, the numerical stability of these models may be compromised when the simulated cells undergo large deformations. We review the features of available 3D DCMs in Table 1.

**Table 1:**
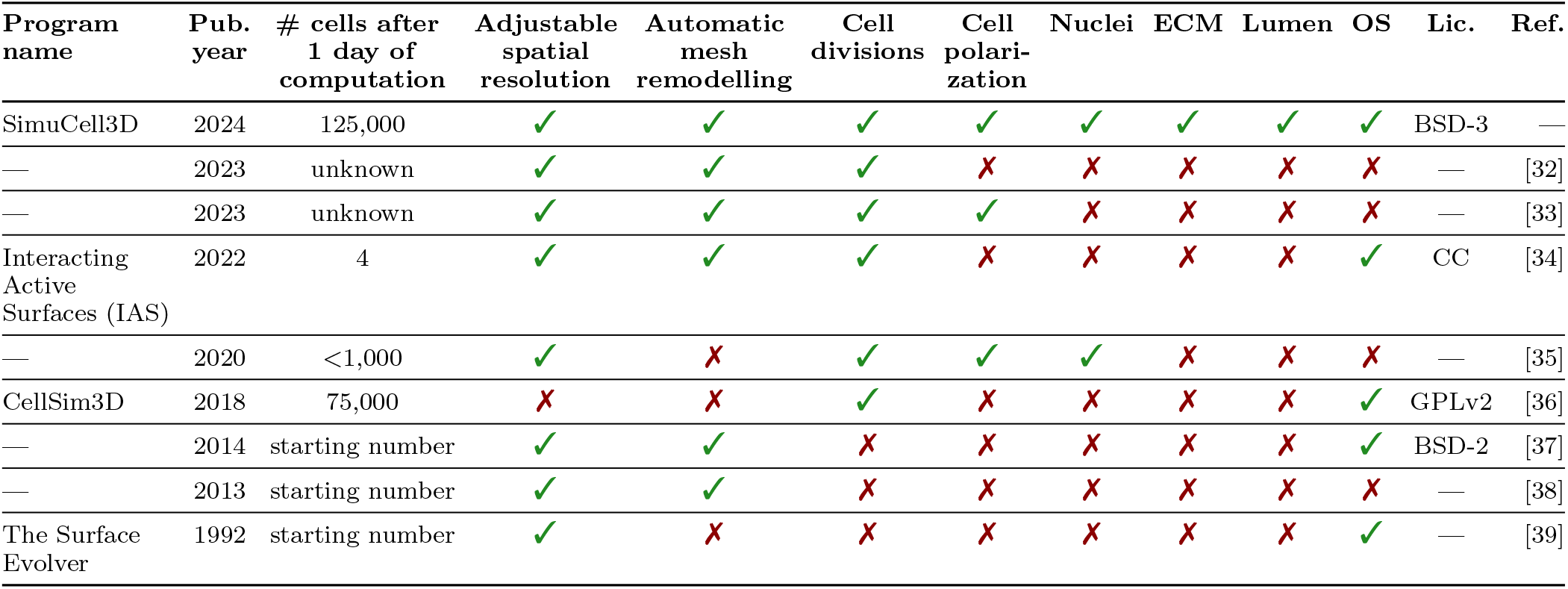
Comparison of SimuCell3D to existing 3D DCM models. Cell numbers are approximate. Pub: Publication. Lic: License. OS: Open source. Ref: References. CC: Creative Commons. GPL: GNU General Public License. BSD: Berkeley Software Distribution.

| Program name | Pub. year | # cells after 1 day of computation | Adjustable spatial resolution | Automatic mesh remodelling | Cell divisions | Cell polarization | Nuclei | ECM | Lumen | OS | Lic. | Ref. |
| --- | --- | --- | --- | --- | --- | --- | --- | --- | --- | --- | --- | --- |
| SimuCell3D | 2024 | 125,000 | ✓ | ✓ | ✓ | ✓ | ✓ | ✓ | ✓ | ✓ | BSD-3 | — |
| — | 2023 | unknown | ✓ | ✓ | ✓ | ✗ | ✗ | ✗ | ✗ | ✗ | — | [32] |
| — | 2023 | unknown | ✓ | ✓ | ✓ | ✓ | ✗ | ✗ | ✗ | ✗ | — | [33] |
| Interacting Active Surfaces (IAS) | 2022 | 4 | ✓ | ✓ | ✓ | ✗ | ✗ | ✗ | ✗ | ✓ | CC | [34] |
| — | 2020 | <1,000 | ✓ | ✗ | ✓ | ✓ | ✓ | ✗ | ✗ | ✗ | — | [35] |
| CellSim3D | 2018 | 75,000 | ✗ | ✗ | ✓ | ✗ | ✗ | ✗ | ✗ | ✓ | GPLv2 | [36] |
| — | 2014 | starting number | ✓ | ✓ | ✗ | ✗ | ✗ | ✗ | ✗ | ✓ | BSD-2 | [37] |
| — | 2013 | starting number | ✓ | ✓ | ✗ | ✗ | ✗ | ✗ | ✗ | ✗ | — | [38] |
| The Surface Evolver | 1992 | starting number | ✓ | ✗ | ✗ | ✗ | ✗ | ✗ | ✗ | ✓ | — | [39] |

Here we present SimuCell3D, a highly efficient open-source DCM in 3D. Thanks to its efficient design, SimuCell3D can simulate tissues composed of dozens of thousands of cells with high spatial resolution. SimuCell3D overcomes the classical trade-off that has so far constrained cell-based models between their resolutions and the number of cells they can simulate. In addition, our program natively allows one to represent intra- and extra-cellular entities such as nuclei, lumens, ECM, and non-uniform mechanical cell membrane properties, as found in polarized cells (Fig. 1a). By combining speed and versatility, SimuCell3D can simulate processes that were not yet amenable to existing numerical methods. We first present the strategy used to represent the biomechanical and geometrical state of cells in SimuCell3D. Then, we demonstrate the computational efficiency of our program and its versatility by simulating various tissue topologies. Given the biological importance of cell polarity, we then present a ray casting method allowing SimuCell3D to automatically polarize cells based on local tissue topology. Finally, we apply SimuCell3D to the study of two morphogenetic problems, the formation and maintenance of layered epithelia, and the cellular organization in a pseudostratified epithelium.

**Figure 1:**
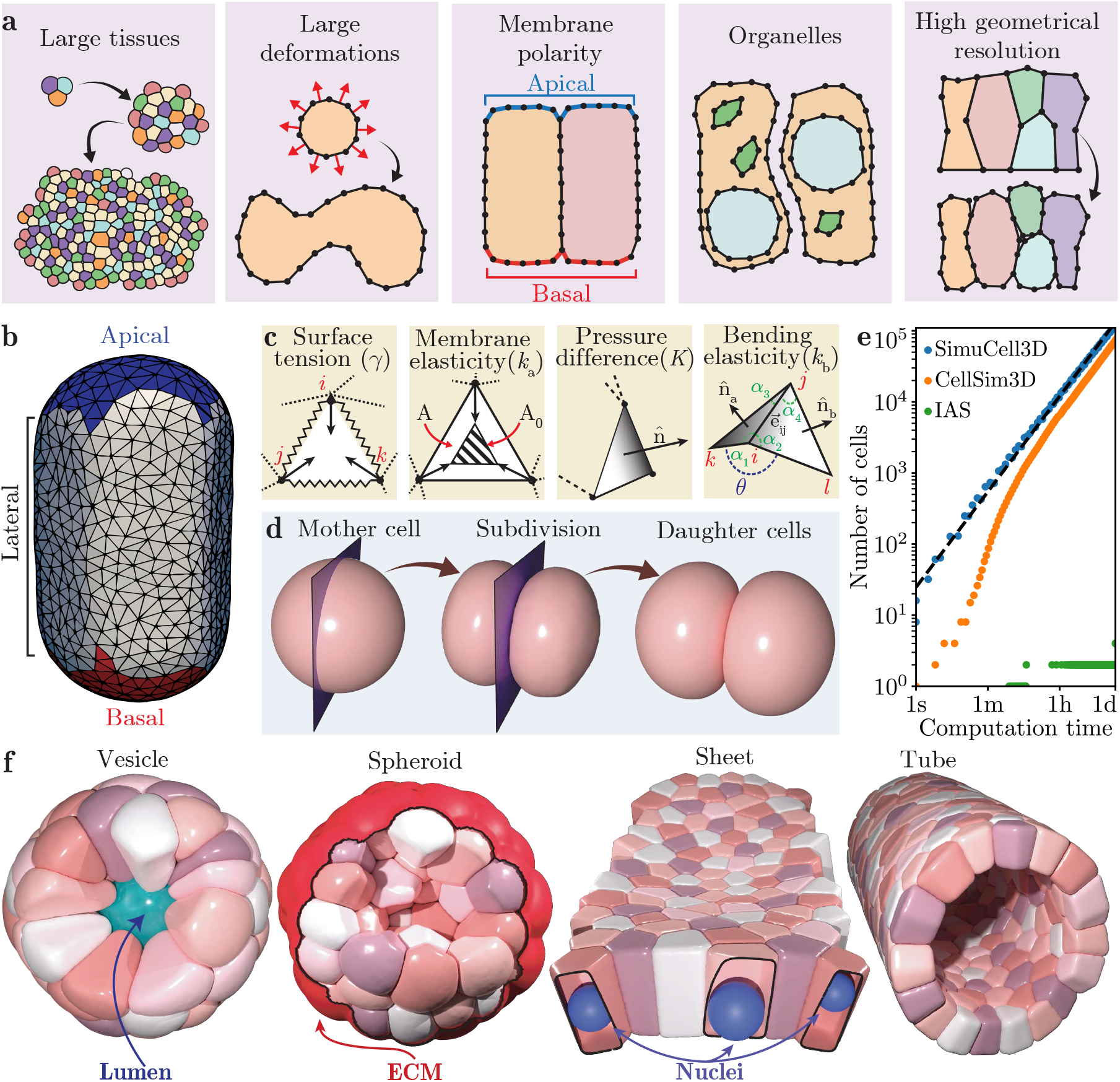
Representation of cellular tissues in SimuCell3D. **a** Schematics of the features cell-based models must possess to be applicable to a broad range of morphogenetic problems. **b**, Representation of cell membranes as closed triangulated surfaces with non-uniform material properties based on cell polarity (colors). **c**, Summary of different forces acting on the triangulated cell membranes. **d**, Illustration of cell division perpendicular to a division plane (purple). **e**, Computational efficiency of different 3D cell-based models in an exponential growth scenario. The dashed black line is a fitted power law 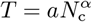 with coefficients *a* = 0.013 s and α = 1.33. **f**, Illustration of different tissue topologies and intra- or extracellular features that can be simulated with SimuCell3D. All surfaces can be non-convex.

## Results

### Biophysical model

SimuCell3D aims to simulate the morphodynamics of cellular tissues at a high spatial resolution with full account of complex cell shapes. The shapes and motion of the simulated cells are not constrained by the model representation, and their mechanical properties are based on the physical principles governing the dynamics of their biological counterparts. This unconstrained representation of the cells is achieved by modeling their surfaces with disjoint closed triangulated surfaces (Fig. 1b). The spatial resolution of these surfaces can be tuned by adjusting the size of their triangles. To ensure that the simulations are initialized at the desired resolution, a custom triangulation algorithm has been incorporated into SimuCell3D (Fig. S1), allowing the use of geometries obtained from microscopy images as starting point of the simulations. A local remeshing algorithm (Fig. S2) preserves the mesh resolution and quality even under large cell deformations. Apart from viscous damping as well as repulsive and adhesive cell-cell contacts, the biomechanical state of each cell membrane is defined by the following energy potential (Fig. 1c):

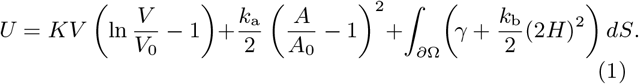

The first term is the energy associated with a net internal pressure, *p = dW/dV =* −*K ln(V/V*_0_ ), which arises from the volumetric strain of the cell cytoplasm, modeled as a slightly compressible fluid. *W* denotes work, *V* and *V*_0_ are the current and target cell volumes, and *K* the cytoplasmic bulk modulus. Shrinkage or growth of cells can be achieved by evolving their target volumes in time. The second energy term allows each cell to actively regulate its membrane area *A* by penalizing deviations from a target value *A*_0_ with an effective isotropic membrane elasticity parameter *k*a. The first term in the surface integral, which runs over the cell surface *∂*Ω, models the tension generated by the cell actomyosin cortex. γ is the isotropic cortical tension, analogous to surface tension of fluid interfaces. The second integrand models the resistance of the cell cortex to bending [68], with H denoting the local mean curvature of the cell membrane, and *k*_b_ its bending rigidity. γ and *k*_b_ are field parameters that can vary along the cell surface according to cell polarity (Fig. 1b).

SimuCell3D offers two distinct contact models to simulate intercellular interactions. The first model mediates interactions through local elastic contact forces, taking into account cell-cell adhesion and volumetric exclusion (Methods, Eq. 2, Fig. S3). Its two constitutive parameters, the adhesion strength ω and the repulsion strength ξ, are field quantities that can vary among cells or membrane regions. This contact model is somewhat mesh resolution-dependent (Fig. S4), just like adhesion in biology will depend on the adhesion protein density. The second contact model mechanically couples the nodes of adjacent cells and directly transfers forces generated on one cell surface to that of the neighboring cell. We validated this second model by reproducing the Young–Dupré relationship in cell doublets and triplets (Fig. S5a,c). The resulting contact mechanics are mesh resolution-independent (Fig. S5a). All parameters related to intercellular interactions are summarized in Table 2.

**Table 2:** Model parameters. Default parameter values are given for the two types of equations of motion implemented in SimuCell3D (dynamic *versus* overdamped). In the parameter dimension, M represents mass, L length, T time. Default values produce a typical tissue growth scenario.

| Symbol | Default<br>dynamic | Default<br>overdamped | Measured | Unit | Dimension | Description | Ref. |
| --- | --- | --- | --- | --- | --- | --- | --- |
| **Cell volume parameters** |  |  |  |  |  |  |  |
| $\rho$ | 1000 | 1000 | 1045 – 1099 | kg/m<sup>3</sup> | M/L<sup>3</sup> | Mass density | [55] |
| $K$ | 2500 | 2500 | 2250 | Pa | M/LT<sup>2</sup> | Bulk modulus | [56] |
| $p_{\max}$ | 2500 | 2500 | 300 – 2200 | Pa | M/LT<sup>2</sup> | Max. internal net pressure | [57–59] |
| $V_{\min}$ | $3.7 \times 10^{-16}$ | $3.7 \times 10^{-16}$ | $2.5 - 3.7 \times 10^{-16}$ | m<sup>3</sup> | L<sup>3</sup> | Min. volume (apoptosis) | [60, 61] |
| $V_{\max}$ | $1.4 \times 10^{-15}$ | $1.4 \times 10^{-15}$ | $0.9 - 1.3 \times 10^{-15}$ | m<sup>3</sup> | L<sup>3</sup> | Max. volume (cell division) | [60, 61] |
| $g$ | $10^{-11}$ | $10^{-11}$ | $0.1 - 1.8 \times 10^{-20}$ | m<sup>3</sup>/s | L<sup>3</sup>/T | Volumetric growth rate | [62, 63] |
| **Cell surface parameters** |  |  |  |  |  |  |  |
| $\gamma$ | 0.001 | 0.001 | 0.0005 – 0.0025 | N/m | M/T<sup>2</sup> | Surface tension | [64–66] |
| $k_b$ | $2 \times 10^{-18}$ | $2 \times 10^{-18}$ | $1 - 2 \times 10^{-18}$ | J | ML<sup>2</sup>/T<sup>2</sup> | Bending stiffness | [67] |
| $k_a$ | $10^{-15}$ | $10^{-15}$ | — | J | ML<sup>2</sup>/T<sup>2</sup> | Area elasticity modulus | — |
| $Q_0$ | 250 | 250 | 300 | — | — | Target isoparametric ratio | [61] |
| $\xi$ | $10^9$ | $10^9$ | — | Pa/m | M/L<sup>2</sup>T<sup>2</sup> | Repulsion strength | — |
| $\omega$ | $10^9$ | $10^9$ | — | Pa/m | M/L<sup>2</sup>T<sup>2</sup> | Adhesion strength | — |
| $H_{\max}$ | $5 \times 10^6$ | $5 \times 10^6$ | — | 1/m | 1/L | Max. coupling curvature | — |
| **Numerical parameters** |  |  |  |  |  |  |  |
| $l_{\min}$ | $2 \times 10^{-7}$ | $2 \times 10^{-7}$ | — | m | L | Minimum edge length | — |
| $c$ | $2 \times 10^{-7}$ | $2 \times 10^{-7}$ | — | m | L | Contact cutoff distance | — |
| $\zeta$ | $2.5 \times 10^{-10}$ | $3 \times 10^{-9}$ | — | kg/s | M/T | Viscous damping coefficient | — |
| $\Delta t$ | $10^{-7}$ | $10^{-7}$ | — | s | T | Time step | — |

SimuCell3D can simulate entities such as nuclei, lumens, and extracellular matrices (ECM) by also representing them with closed triangulated surfaces like the cell membranes. To model cell death, cells can be removed from the tissue if their volume drops below a minimum threshold *V*_min_. Conversely, if cell volumes exceed the maximal value *V*_max_, they undergo cytokinesis (Fig. 1d). The division plane can be randomly oriented or perpendicular to the longest cell axis (Hertwig’s rule). A cell division only takes a few microseconds of computation time, allowing the simulation of tissues with high cell division rates. To demonstrate the computational efficiency and stability of our program, we simulated the exponential growth of a tissue in an out-of-equilibrium regime with the growth rate pushed to the limit, starting from a single cell (Fig. 1e, Movie S1). The cells in this test are simulated without nucleus. Only one day of computation time is required to grow the tissue to 125,000 cells on an Intel Xeon W-2245 CPU (8 cores, 3.9 GHz) using 16 threads, for cells that possess 121 nodes and 238 triangular faces on average. The total time complexity of such a simulation is 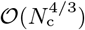, where Nc is the number of cells in the tissue, which is equivalent to the scaling observed in 2D simulations [51]. Under similar settings, we tested the performance of CellSim3D [36] and Interacting Active Surfaces (IAS) [34], two other cell-based 3D models offering low and high spatial resolution, respectively. CellSim3D generated a tissue of 75,000 cells in a day of computation time while IAS produced a tissue of 4 cells in the same amount of time. CellSim3D achieves comparable performance as our program by constraining the cell geometries to simple spherical shapes with a fixed number of nodes. SimuCell3D thus offers the performance of low-resolution models such as CellSim3D while possessing the flexibility and accuracy of high-resolution models like IAS. SimuCell3D is parallelized with OpenMP. The parallelization efficiency follows Amdahl’s law (Fig. S6). To showcase the versatility of our program, we simulated various tissue topologies such as a vesicle, a bulk spheroid, a sheet, and a tube, alongside several intra- or extracellular built-in features like lumens, nuclei, and ECM (Fig. 1f, Movie S2).

### Cell membrane polarization

Cells form regions with distinct biochemical and mechanical properties along their cytoplasmic membranes. Correct establishment of this cell polarity is crucial to numerous developmental processes [69]. Its impairment has also been linked to the onset of tumor formation [70]. SimuCell3D takes cell polarity into account by allowing the triangular faces to be of different types with distinct mechanical parameters γ, *k*_b_, ω, and ξ. Two mechanisms are implemented to automatically identify different regions on the cell surfaces. In the first, lateral sides are inferred from the face contact information, leaving regions that are not in contact as either apical or basal. The second, more robust and versatile algorithm is based on a spatial partitioning of the simulation domain into voxels representing one of four possible regions: cell boundaries, luminal, cytoplasmic, and external (Fig. 2a-c). Voxels containing mesh nodes are marked as boundary voxels. The remaining unmarked voxels are clustered with the Hoshen–Kopelman algorithm [71]. The different voxel clusters thus created are then labeled as cytoplasmic, luminal, or external based on their positions in the discretized simulation space. Then, each surface triangle probes its environment by casting a ray in the direction of its outward normal to detect which type of region it faces (Fig. 2d). The type of voxel the ray first passes through determines whether the mesh triangle is lateral (facing another cell), apical (facing an enclosed volume such as a lumen) or basal (facing the surrounding medium or ECM). Iteration over all mesh triangles thus tags the entire surface (Fig. 2e). We demonstrate the strength of this approach by reproducing *in silico* a monolayer prostate organoid whose cells exhibit apico-basal polarity (Fig. 2f). The cell surfaces were extracted from 3D microscopy images with CellPose [72] (Fig. 2g). SimuCell3D then reproduced the organoid with correct tissue polarity (Fig. 2h) without requiring any input on tissue orientation or topology by the user.

**Figure 2:**
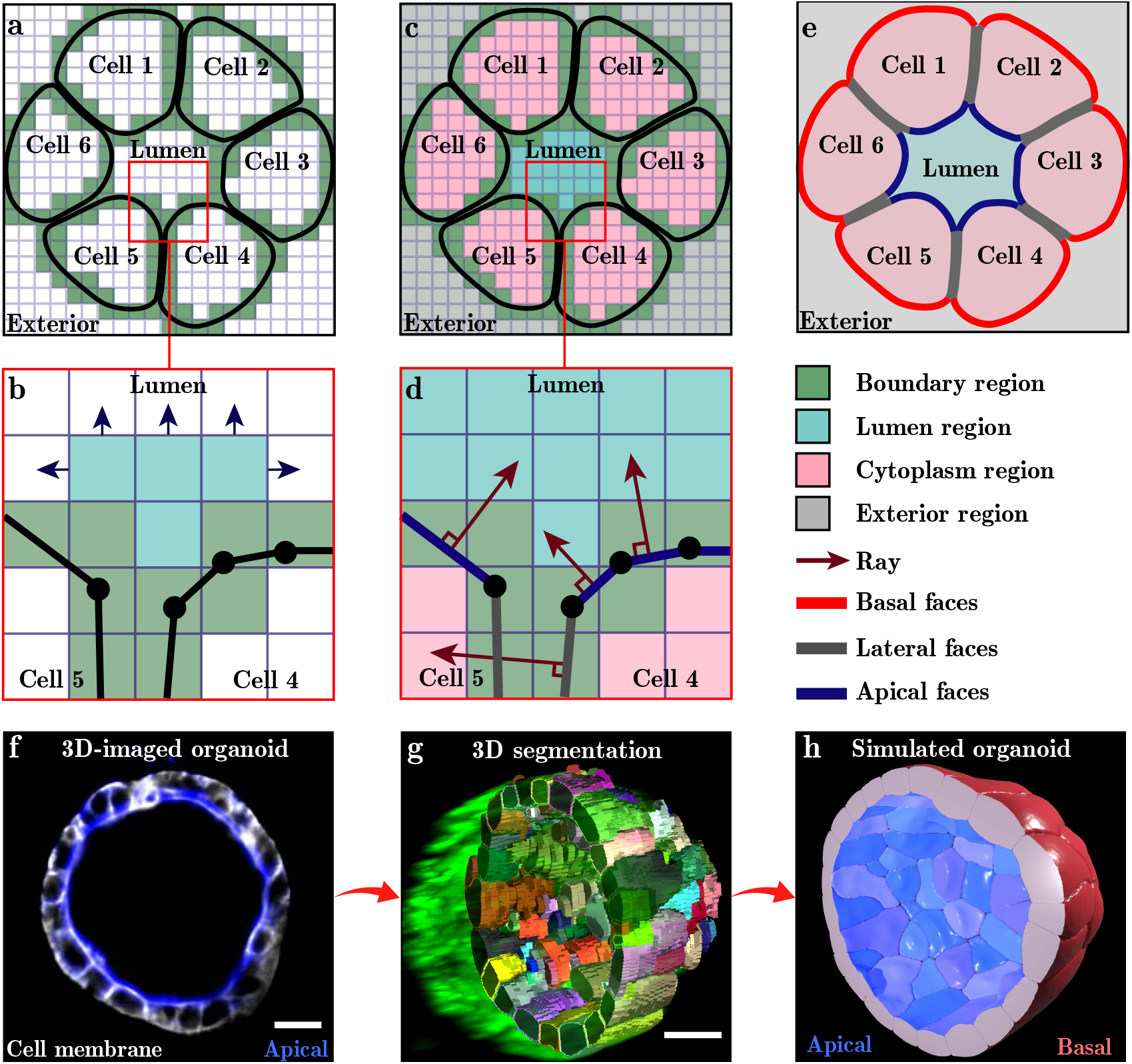
Automatic cell surface polarization algorithm. For visual clarity, the process is shown in 2D. **a**, The simulation space is discretized with uniformly sized cubic voxels. All voxels that intersect with the cell surfaces are marked as boundary voxels (green). **b**, All remaining voxels are clustered with the Hoshen–Kopelman algorithm. **c**, Voxel clusters are tagged as lumen (turquoise), cytoplasm (pink) or exterior (gray). **d**, A ray is cast from the center of each face in outward normal direction. The first voxel (other than a boundary) that it intersects with indicates with which region the face is in contact. **e**, Facial types (red, gray, blue) are assigned based on the regions with which they are in contact. **f**, Cross-sectional light-sheet microscopy image of a mouse prostate organoid. Cell polarity is visualized by Ezrin staining (apical side, blue). **g**, 3D cell segmentation of the organoid. **h**, Simulated organoid with automatically polarized epithelial surfaces (blue, red). The different membrane regions can possess different surface tensions, bending rigidities, adhesion strengths and repulsion strengths. Scale bars represent 15 μm.

## Application 1: Transition from monolayer to multilayer tissue

We now demonstrate how SimuCell3D can be used to gain insight into the cellular dynamics of biological tissues. As a first showcase, we investigate the relationship between biomechanical cell parameters and the internal structure of a tissue as a mono- or multilayer. Such a difference in cellular organization is particularly striking between different types of epithelial tissues [73]. Strong evidence suggests that this variability is the result of an interplay between intra-cellular surface tension and inter-cellular adhesion [74, 75]. In a tug of war with cortical tension, in which the actomyosin cortex tends to minimize the cell surface area, adhesion molecules between adjacent cells tend to increase it. We investigated this competition by numerically exploring the parameter space spanned by adhesion strength and surface tension. The simulations were initialized with a spherical monolayer vesicle consisting of 432 columnar cells generated from a Voronoi tesselation of the sphere (Fig. 3a). All cells were initially in contact on their apical sides with a luminal region and on their basal sides with an ECM encasing the tissue. Note that no ECM located at the apical side of the cells nor any adhesion belt were considered in these simulations. The cells were grown at a uniform volumetric rate without division until they had doubled in size, while the luminal target volume was preserved. Despite cellular rearrangements caused by growth, we observed the maintenance of the monolayer structure in simulations with low surface tension (Fig. 3b). Strong cortical tension, on the other hand, leads to stratification (Fig. 3c). We quantified the resulting number of cell layers by converting the tissue into a graph representing cell connectivity and computing the shortest path percolating from the lumen to the ECM (Fig. 3d). Parameter values were non-dimensionalized with *l* = ⟨*V* (t = 0)⟩^1*/*3^ as a characteristic length scale, and the cytoplasmic bulk modulus *K* as a characteristic energy density. Our exploration of the parameter space revealed that, under the prescribed conditions, the layering of the tissue is essentially regulated by the tension of the actomyosin cortex alone (Fig. 3e). An increase in the normalized surface tension 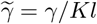, from 0.02 to 0.10 was sufficient to break the monolayer arrangement and force the tissue into a stratified structure. Conversely, an increase by two orders of magnitude in the normalized adhesion strength 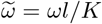, between the cells did not disrupt the monolayer integrity. As the cells lose their apico-basal connectivity at stronger surface tension, they adopt a more spherical shape that minimizes their surface area, as measured by their sphericity Ψ = π^1*/*3^(6*V*)^2*/*3^/*A* (Fig. 3f). These simulations highlight the potential of SimuCell3D to quantitatively address open questions in tissue development and cancer progression, the latter being linked to a loss of structural tissue integrity on the cellular level [76].

**Figure 3:**
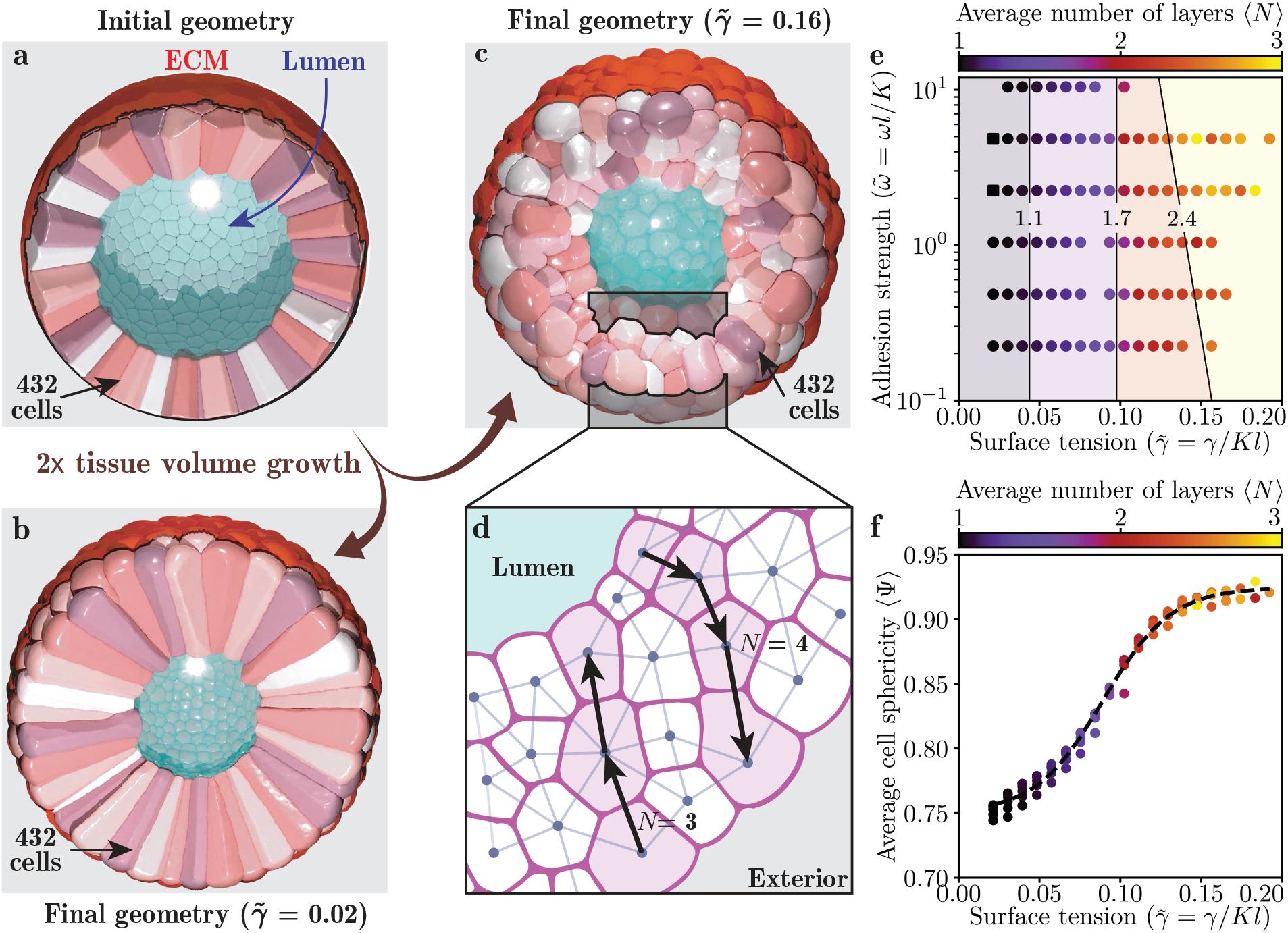
Impact of biomechanical cell properties on tissue structure. **a**, Initial tissue geometry, a hollow spherical vesicle made up of 432 columnar epithelial cells. The central luminal region (turquoise) was represented by a non-growing volume. On its basal side, the tissue was encased by a surface representing the ECM (red). **b**, Final monolayer conformation after epithelial volume doubling with weak cortical tension. **c**, Final multilayer conformation at strong cortical tension. **d**, Schematic cross section through the epithelium with a cell connectivity graph on which the layer number N was determined. **e**, Impact of cell surface tension and adhesion strength on the average number of cell layers. Isolines represent support vector machine discriminants. Each data point corresponds to the final state of one numerical simulation. Simulations in which the whole tissue remained a monolayer (N ≡ 1) are shown as squares. **f**, Effect of cortical tension on cell shape, as measured by their sphericity, and on the number of layers (colors). The dashed black curve is a fitted logistic function.

### Application 2: Formation and maintenance of pseudostratification in epithelia

Pseudostratified epithelia are single-layered epithelia that are easily mistaken as stratified when analysed in 2D sections because of the dispersion of their nuclei along the apical-basal axis [61]. Their ubiquity across different species during development [77] suggests that the pseudostratified structure can confer an advantage over simpler cellular arrangements, possibly linked to patterning precision [78]. How this structure is acquired and maintained under growth and morphogenetic deformation is still largely unknown. In this second case study, we demonstrate how SimuCell3D may be used to gain mechanistic insight into the elusive pseudostratification process. We initialized simulations with a patch of 70 cells segmented from light-sheet microscopy images of the developing pseudostratified mouse lung epithelium [61] (Fig. 4a). Among these 70 cells, the 21 interior cells were allowed to move freely while the rest on the periphery of the patch acted as static boundaries. The simulated cells all contained a nucleus (Fig. 4b, blue) and did neither grow nor divide during the simulations, but deformed to minimize their potential energy, until static equilibrium was reached. We again examined the interplay between cell surface tension 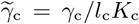, subscript “c” for cell) and adhesion strength 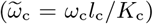 (Fig. 4c,d). The normalized surface tension of the nuclei 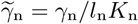, subscript “n” for nucleus) was kept constant at 0.24 in these simulations, and they were non-adhesive (ω_n_ = 0). In the explored region of the parameter space, we observed two morphological cell regimes (I and II) with a continuous transition in between, along which an intermediate physiological range can be identified (Fig. 4c). In regime I, the cell shape is dominated by the effect of surface tension. Some of the cells segregated in response to the strong surface area minimization tendency (Fig. 4c, left), facilitated by weak lateral adhesion. Cells in this regime reduced their lateral cell-cell contact area fraction ϕ (Fig. 4d) and also possessed fewer neighbors, as measured by their coordination number z (Fig. 4e). In regime II, the effect of adhesion dominates over surface tension, allowing neighboring cells to maximize their mutual contact areas (Fig. 4c, right; Fig. 4d) as well as their coordination number (Fig. 4e). In between these extremes, a balance between adhesion strength and cortical tension yields physiological cell shapes corresponding to those imaged (Fig. 4c, middle). This morphological similarity can be exploited to infer the mechanical properties of *in vivo* pseudo-stratified cells (Fig. 4d,e). Moreover, besides informing on the mechanical state of cells, SimuCell3D unveiled in this second case study the possibility that pseudostratified tissues could be formed from cells with identical mechanical properties.

**Figure 4:**
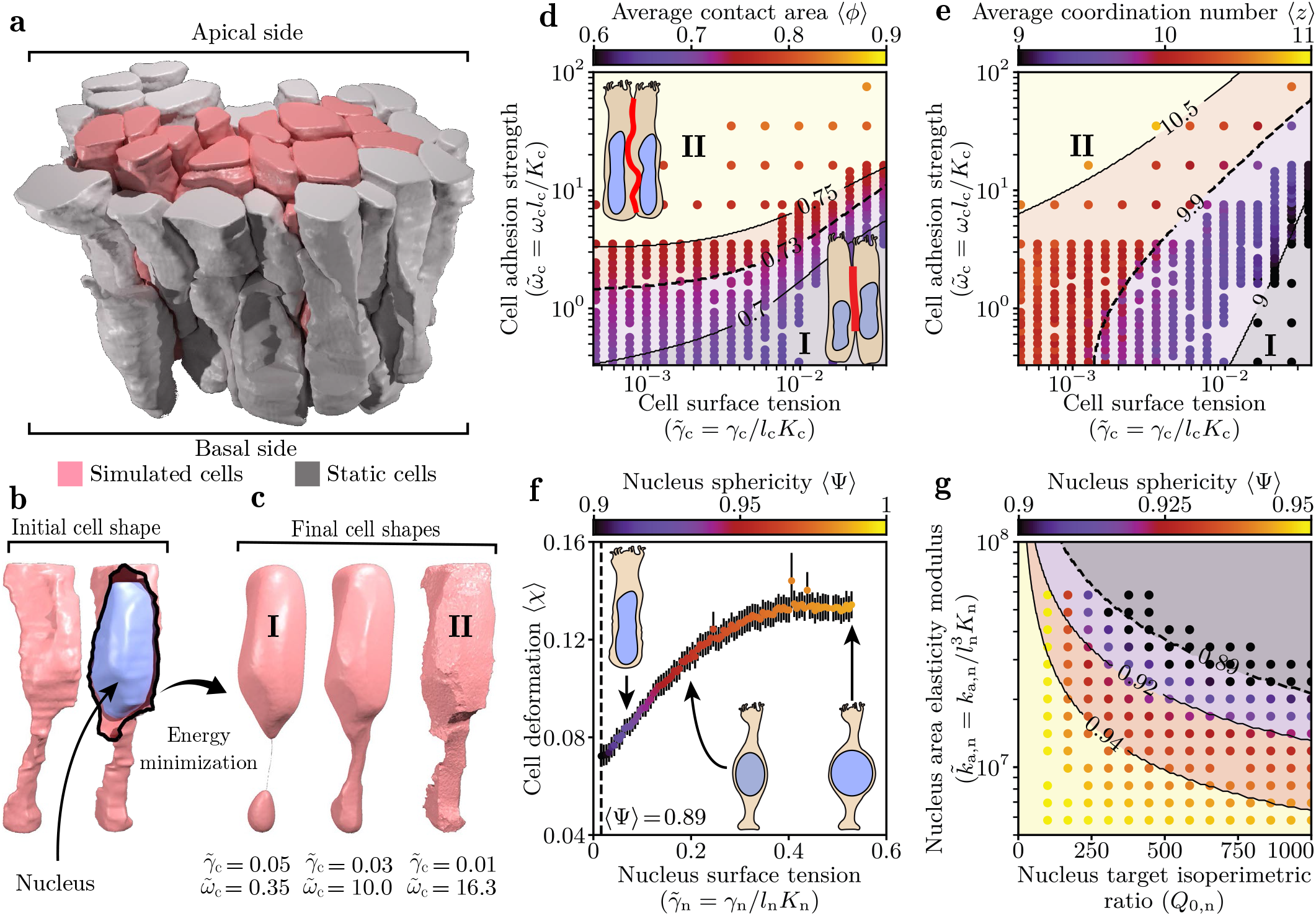
Simulation of 3D cell organization pseudostratified epithelia. **a**, Initial geometry imported into SimuCell3D, a patch of 70 cells from the developing pseudostratified mouse lung epithelium, segmented and triangulated from light-sheet imaging [61]. 49 gray cells act as rigid boundaries for the 21 simulated pink cells. **b**, All cells contain a nucleus (blue). **c**, Based on the cell surface tension and adhesion strength, the cells adopt different shapes (regimes I and II). **d**, Mean fraction of cell surface area in contact with other cells (red line in illustration), as a function of the adhesion strength and cortical tension. **e**, Mean cell coordination number in the same parameter space. **f**, Cell deformation as a function of the nuclear surface tension. Error bars represent the standard error. **g**, Average nuclear sphericity as a function of the nucleus area elasticity modulus and target isoperimetric ratio. Isolines in d,e,g represent support vector machine discriminants. Dashed curves in d,e,f, and g represent the physiological values measured from the segmented geometries. Each data point corresponds to the equilibrium state obtained from one numerical simulation.

Subsequently, we used SimuCell3D to investigate the effect of mechanical properties of the nuclei on the pseudostratified cell organization (Fig. 4f,g). In these simulations, the cell surface tension 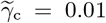, and adhesion strength 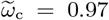, were fixed. By varying the nuclear surface tension 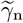, we were able to create nuclei rigid enough to deform the cell membranes (Fig. 4f). Cell deformation was measured by comparing the equilibrium cell shape in the presence of a nucleus *versus* in its absence, quantified by the intersection over union: χ = 1 − IoU(Ω with nucleus, Ω without nucleus). We observe an increase of the average cell deformation ⟨χ⟩ with the nucleus surface tension 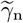 until the nuclei obtain spherical shapes at 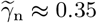. It then saturates at ⟨χ⟩ ≈ 0.13, as nuclear tension increases further. The average sphericity of the nuclei has been measured in the segmented geometries at 0.89, suggesting a low cortical stiffness of the nuclear envelopes relative to the cytoplasmic membranes.

SimuCell3D also allows one to directly modulate the shapes of nuclei or cells by concurrently varying their target isoperimetric ratios 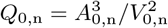, and area elasticity moduli *k*_a,n_ (Fig. 4g). Simultaneously increasing Q_0,n_ and *k*_a,n_ drives the equilibrium shapes of nuclei away from a sphere. Conversely, nuclei with small values of Q_0,n_ and *k*_a,n_ possess more spherical shapes. The ability to thus change the stiffness or shape of the nuclei opens up opportunities to study the dynamics of interkinetic nuclear migration [79].

## Discussion

In this study, we presented SimuCell3D, a three-dimensional cell-based model that permits the simulation of very large, growing, and deforming tissues at high cellular and subcellular resolution with apical-basal polarity, ECM, and luminal spaces. SimuCell3D achieves this through an extensive level of parameterization and a built-in polarization algorithm. Moreover, due to its efficient implementation, SimuCell3D permits the simulation of phenomena involving tens of thousands of cells at a high resolution for the first time.

This now permits the in-depth *in silico* investigation of the mechanical properties and behavior of cells to understand the mechanisms that regulate tissue homeostasis and morphogenesis. While the current simulations were carried out with linear mechanical models, non-linear material behavior (visco-elasticity, hyperelasticity) could readily be implemented to study their effect on morphogenesis. Moreover, besides nuclei, organelles and endocytosis could be easily represented. As such, processes such as interkinetic nuclear migration in pseudostratified epithelia could be simulated at unprecedented resolution to address open questions regarding the driving forces.

As we showed, SimuCell3D can be used to predict the global tissue morphologies that emerge from individual mechanical cell properties. Specifically, when the morphological features of the tissues are known, SimuCell3D can be used to infer the region of the mechanical parameter space in which the cells are located. Our exploration of the cellular parameter space in this study was mainly limited to the subspace spanned by cell cortical tension and adhesion strength. This subspace is insufficient to reproduce the wealth of morphogenetic events observed *in vivo*.

In other contexts, exploration of higher dimensional parameter spaces will undoubtedly be necessary. In these circumstances, SimuCell3D can be coupled with gradient-free parameter estimation techniques to accurately infer the cell properties that lead to the measured morphological tissue features.

SimuCell3D is readily extendable to accommodate more features in the future. Relevant possible extensions include subcellular components such as adhesion belts, frictional forces which play an important role in the morphogenesis of some tissues [80], as implemented in pre-existing models [33, 35], tension fluctuations [81], and reaction-diffusion models to couple the biomechanical tissue model with chemical signalling. In this way, chemical and mechanical symmetry-breaking mechanisms could be combined and their effects could be simulated at cellular resolution. Finally, the cell-based simulations could be combined with continuum models to simulate the behaviour of larger tissues at varying resolution, and to derive adequate material models for the continuum description from cell-based simulations.

## Methods

### Mesh operations

#### Local mesh adaptation

SimuCell3D geometrically represents cells by closed triangulated surfaces whose edge lengths *l* are maintained within the range [*l*_min_, *l*_max_] with a local mesh adaptation method. The minimum edge length *l*_min_ is a model parameter, whereas *l*_max_ = 3*l*_min_, a value that works well in most practical applications, is automatically set. When the length of an edge exceeds *l*_max_, the local mesh adaptation method splits it in two (Fig. S2a), adding one node and two faces to the mesh. The two nodes constituting the divided edge transfer a third of their momentum to the newly created middle node to ensure momentum continuity. When an edge shrinks to a length below *l*min, it is collapsed into a node whose new momentum is the sum of the merged nodes (Fig. S2b). This merging process eliminates one node and two faces from the cell mesh. To prevent triangles with vanishing area, this operation is allowed only when the two nodes to be merged share exactly two other nodes among those connected to them through edges.

Triangular faces with high isoperimetric ratios can be a source of numerical instability. An edge swap operation prevents their formation. First, the quality score 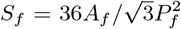 of each face *f* is computed, where P_*f*_ is its perimeter and *A*_*f*_ its area. Undesirable faces with high isoperimetric ratios have scores tending to zero, whereas S_*f*_ = 1 for equilateral triangles. Faces with a score S_*f*_ < 0.3 are eliminated by an edge swap operation (Fig. S2c) that locally reconnects mesh nodes, but leaves them otherwise unaffected.

#### Initial triangulation

A flexible triangulation algorithm ensures that simulations are initialized with meshes that respect the edge length bounds (Fig. S1a). The procedure takes an initial geometry of the tissue as input, with cell meshes that are not necessarily triangular yet, in the widely used VTK format [82]. The cell surfaces are then individually sampled with the Poisson Disk Sampling algorithm [83] (Fig. S1b) with a minimal point separation of *l*_min_. The Ball Pivoting Algorithm [84, 85] then separately re-triangulates the surface of each cell based on its Poisson point cloud (Fig. S1c). The resulting meshes have edge lengths *l* ≥ *l*_min_, rarely exceeding *l*max. Edges with lengths *l* > *l*_max_ are removed before the simulation starts with the edge division operation described above.

#### Cell division

Cells are divided based on a volume threshold, *i*.*e*., if *V* > *V*_max_. They are bisected by a plane running through their centroid, whose orientation can depend on the cell type. The orientation is either random or perpendicular to the cell’s longest axis, as given by the eigenvector belonging to the smallest eigenvalue of its covariance matrix. During division, the cutting planes are re-triangulated in a manner respecting the edge length bounds. On the untriangulated region of the daughter cells, points are first sampled with the Poisson Point Cloud algorithm [83], which are then connected with the 2D Delaunay triangulation algorithm. This method avoids a retriangulation of the parts of the cell surface inherited from the mother cell.

### Cell volume and area calculation

The cell volume is calculated with a three-dimensional variant of the shoelace formula:

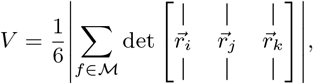

where 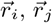, and 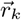 are the nodal positions of face f (Fig. 1c). The summation runs over all the triangular faces of the cell mesh ℳ. The cell surface area is obtained by summing the areas of its faces:

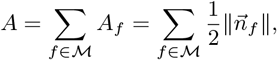

Where 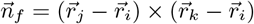, is the unnormalized outward normal of face *f*.

### Time integration

SimuCell3D offers two modes of time propagation, solving either the dynamic or overdamped equations of motion for the nodal positions 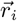,

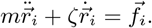

The nodal mass *m* is obtained from the cell volume *V* and mass density *ρ* as m = *ρV* /*N*_*n*_, where *N*_n_ is the total number of nodes in the cell mesh. 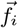 is the nodal force vector (specified below) and ζ the viscous damping coefficient. The first mode resolves elastic oscillations, making it suited for phenomena on short timescales. The nodal positions 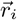 and linear momenta 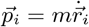 are integrated with the semi-implicit Euler scheme:

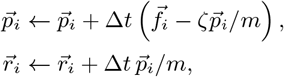

where Δt is a fixed time increment. The second mode neglects inertial effects 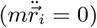 and is therefore suitable for systems dominated by viscous relaxation toward a quasi-static equilibrium. The overdamped equations of motion are solved with the forward Euler scheme:

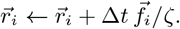

The simulations presented in Figs. 1, 3 and 4 were solved with the dynamic model. Simulation snapshots are written at regular time intervals in VTK format for post-processing and visualization in ParaView (Kitware, Inc.).

### Nodal forces

The total conservative nodal forces 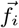 derive from the cell potential energy (Eq. 1) and the intercellular interaction model. They are given by the sum of the surface tension forces, 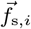, the membrane area elasticity forces, 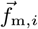, pressure forces exerted by the cytoplasm, 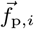, the bending forces, 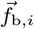, and contact forces due to adhesion and steric repulsion, 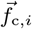:

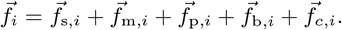

Each of these contributions is detailed in the following paragraphs.

#### Surface tension

The surface tension force is given by the negative gradient of the surface tension energy with respect to the nodal position. Since the position of node i only affects the areas of the set of faces ℱ*i* sharing this node, it is given by

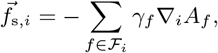

where γ_*f*_ and A_*f*_ are the surface tension and area of face *f*. For triangles with nodes *i, j, k* oriented clockwise (Fig. 1c), the area gradient reads

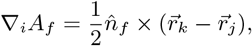

where 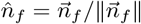, is the normalized face normal vector.

#### Membrane area elasticity

Similarly, the membrane force is obtained by taking the gradient of the cell membrane area energy with respect to 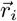:

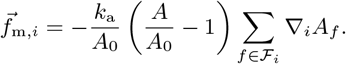

The target area *A*_0_ is coupled to the target volume *V*_0_ via 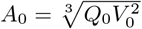, where *Q*_0_ is the target isoperimetric ratio of the cell, which can be set by the user. For a sphere, *Q*_0_ = 36π ≈ 113.

#### Pressure

The cell-internal net pressure generated by the cytoplasm reads

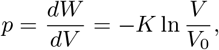

where *W* = −*KV* (ln[*V*/*V*0] − 1), is the work associated with a deviation of the cell volume *V* from its reference value *V*_0_. To model cell growth, *V*_0_ can evolve over time according to prescribed growth laws, such as the linear form d*V*_0_/dt = g, where g is a constant volumetric growth rate that can vary from cell to cell. If desired, the pressure difference between the cell cytoplasm and the external medium can be capped at a predefined threshold *p*_max_, *i*.*e*., *p* ← min{*p, p*_max_}. The pressure force exerted on a subset of the cell surface 𝒮 ⊂ *∂*Ω (where Ω is the cell domain) is given by

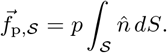

If the subset 𝒮 of the cell surface is planar, like the triangular faces *f* used to discretize the cell geometry, this simplifies to

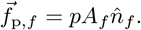

The pressure force applied on each node of the cell mesh therefore follows as

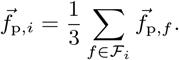

#### Membrane bending

The contribution of bending to the cell potential energy can be approximated with the discrete bending energy [86]

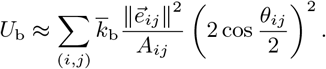

in which the sum runs over all pairs of nodes (*i, j*) of the surface mesh connected by an edge. Each edge connects two faces *a*, b that form a diamond region composed of four nodes *i, j, k, l* (Fig. 1c). 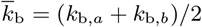 is the average bending stiffness of the faces *a* and b, 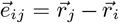 the vector pointing from node i to j, *A*_*ij*_ = *A*_*a*_ + *A*_*b*_ the sum of the two face areas, and *θ*_*ij*_ the dihedral angle between the two faces:

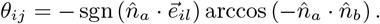

The sign of the dot product between the normal of face *a* 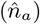 and the vector 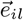 is used to distinguish between concave and convex hinges. The bending forces resulting from this discrete bending energy can be calculated independently for each of the four nodes, *q* ∈ {*i, j, k, l*}, as

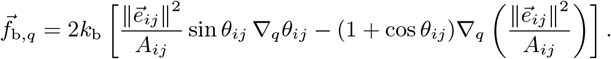

For the gradients ∇_*q*_ θ_*ij*_ and 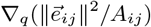 we refer the interested reader to [86]. The total bending force at node *i*, 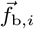, then follows as the sum of bending forces over all diamond regions involving that node.

#### Intercellular contacts

SimuCell3D offers two different contact models that vary in their ways of exchanging contact forces between adjacent cells, but in the current version, it does not take friction into account. (For possible ways to include frictional effects, see e.g. [35, 51].) The first model connects adjacent pairs of faces {*f*_*a*_, *f*_*b*_} with elastic springs, while the second tightly couples pairs of adjacent nodes {*n*_*a*_, *n*_*b*_}.

The spring-based model applies contact forces on pairs of adjacent faces with a magnitude based on the signed distance 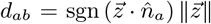 between the two mesh elements, where 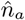 is the unit normal of face *a* and 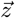 is the shortest vector between the two mesh elements. A contact stress is then calculated with the piece-wise expression

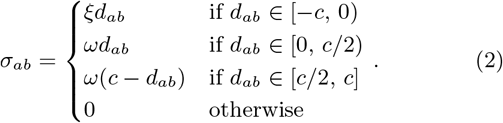

When two neighboring cells interpenetrate, *d*_*ab*_ is negative, and the contact stress is repulsive. ξ is the repulsion coefficient. On the other hand, when *d*_*ab*_ is positive, the contact stress is adhesive. In this regime, the contact model follows a bilinear traction-separation law (Fig. S1). *ω* denotes the adhesion coefficient. The contact stress σ_*ab*_ thus obtained is translated into a force by integrating the contact stress over the contact surface area *A*_*ab*_:

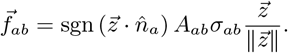

*A*_*ab*_ = min{*A*_*a*_, *A*_*b*_} if the contact forces are computed between pairs of faces {*f*_*a*_, *f*_*b*_}, whereas *A*_*ab*_ = *A*_*a*_ if the contact forces are calculated between pairs of faces and vertices {*f*_*a*_, *v*_*b*_}. In the first case, the force is linearly distributed to the nodes of the face *a*, {*i, j, k*}, and the nodes of the face b, {*l, m, n*}:

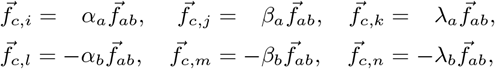

(*α*_*a*_, *β*_*a*_, *λ*_*a*_) and (*α*_*b*_, *β*_*b*_, *λ*_*b*_) are the barycentric coordinates of the closest points of approach on the faces *a* and *b*, respectively.

The second contact model eliminates the need for a finite adhesion strength parameter *ω* by establishing a tight coupling between node pairs {*n*_*a*_, *n*_*b*_} whose distance is smaller than the contact cutoff c. T he t wo n odes a re r elocated t o t heir average location 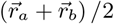, and the forces and momenta acting on each node are transmitted to their partner such that both nodes subsequently follow the same dynamics: 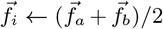 and 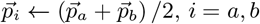. To allow two adjacent cells to detach from each other, node pairs are coupled only if the local mean curvature of both cell surfaces lies below the threshold Hmax (Table 2). Coupled node pairs are redetermined in each timestep, and each node is allowed to be coupled to no more than one other node.

## Acknowledgements

We thank Franziska Lampart for providing the prostate organoid images presented in Fig. 2. This work was partially funded by SNF Sinergia grant CRSII5 170930.

## Author contributions

Concept: RV, DI. Model and algorithm development: SR, RV. Implementation: SR. Numerical simulations: SR. Figures: SR. Writing: SR, RV, DI.

## Code Availability

SimuCell3D is open-source and freely available as a public git repository at https://git.bsse.ethz.ch/iber/ Pub*l*ic*a*tions/2023_runser_3d_ce*ll*_mode*l* from the day of publication. It is released under the 3-clause BSD license.

