## Supplementary Figures for "SimuCell3D: 3D Simulation of Tissue Mechanics with Cell Polarization"

### 3D Simulation of Tissue Mechanics with Cell Polarization: Supplementary Information

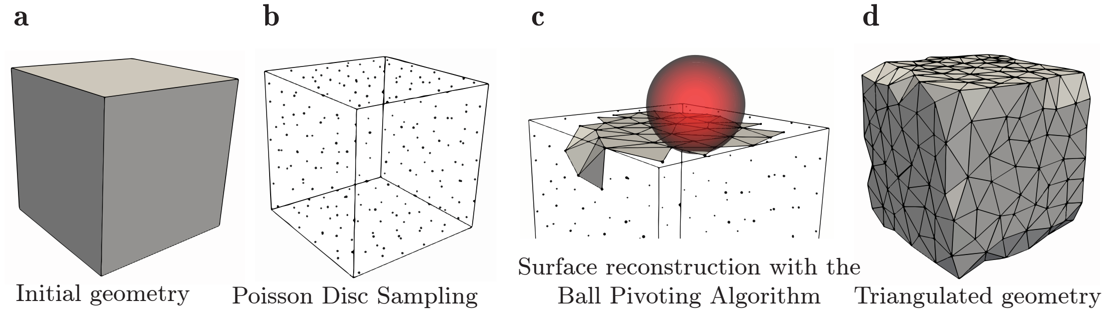


**Figure S1: Initial surface triangulation.** **A**, Arbitrary surface meshes can be loaded by the program, here a cube for illustration. **B**, Point samples are generated on the surface with the Poisson disc sampling method, with a minimal distance of $l_{\min}$ between points. **C**, The Poisson disc point cloud is then used by the Ball Pivoting Algorithm to reconstruct the surface of the initial geometry with triangles. The Ball Pivoting Algorithm connects triplets of points into triangular faces if a ball (red sphere) can simultaneously touch them without containing any other point. **D**, The procedure is terminated when the reconstructed surface is watertight, and the resulting triangulation(s) are used by SimuCell3D to simulate cellular components.

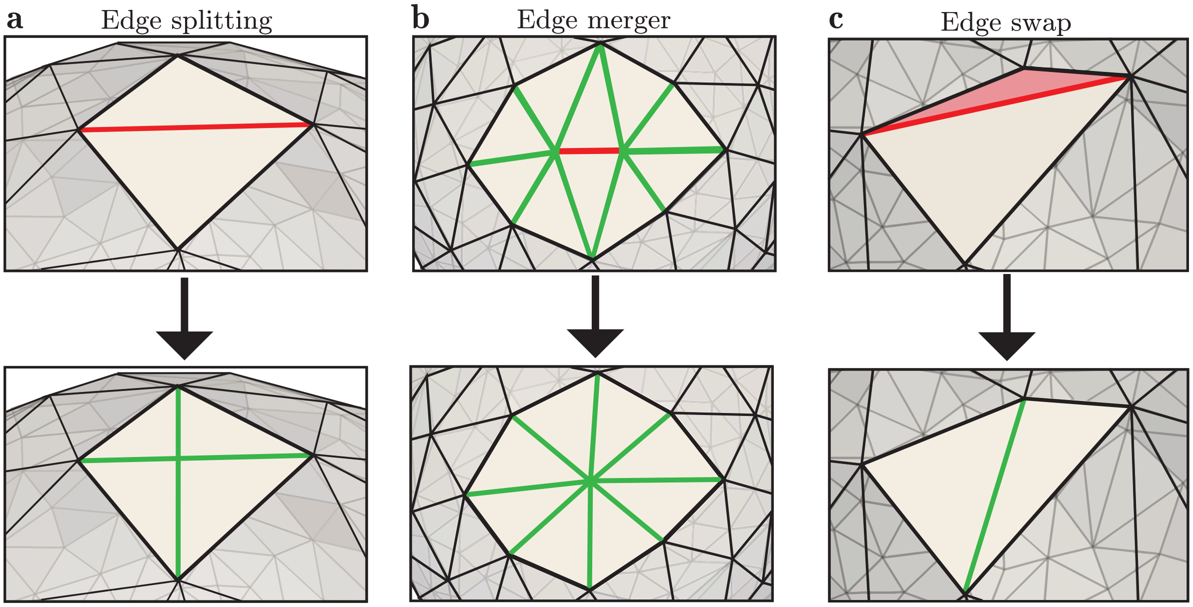


**Figure S2: Local mesh adaptation.** **A**, Edges whose length exceeds $l_{\max}$ (red) are split into two, generating two new triangles and a new node in the middle. **B**, Edges whose length subceeds $l_{\min}$ (red) are collapsed into a node at the center, removing the two triangles sharing it. **C**, Triangles with high isoperimetric ratio (red) are prevented by swapping the orientation of their longest edge.

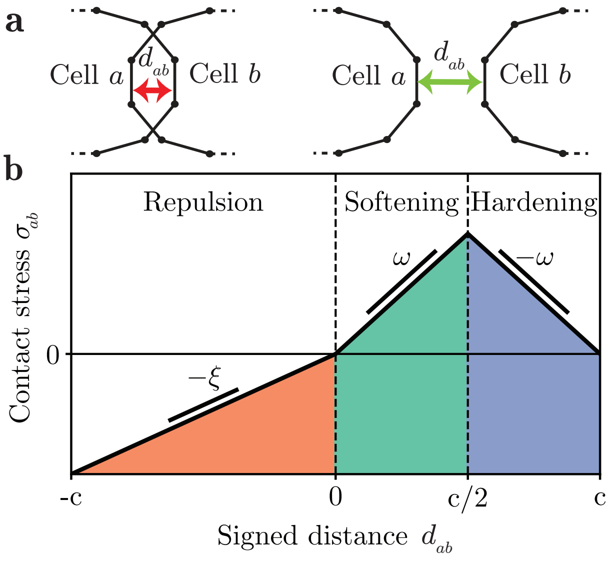


**Figure S3: Cell contact forces. A**, Schematic of two interpenetrating cells (negative distance $d_{ab}$, left) and two adhering cells (positive distance $d_{ab}$, right). **B**, A bilinear adhesive traction-separation law governs the contact mechanics between adjacent cells, symmetrically separated into a hardening regime (green) in which adhesion forces increase linearly with separation, and a softening regime (blue) in which forces decrease linearly, to ensure force continuity at a separation of $d_{ab} = c$. At negative separations, repulsive forces proportional to the penetration depth are exchanged (red). The slopes $\omega$ and $\zeta$ control the rigidity of the intercellular mechanical interactions.

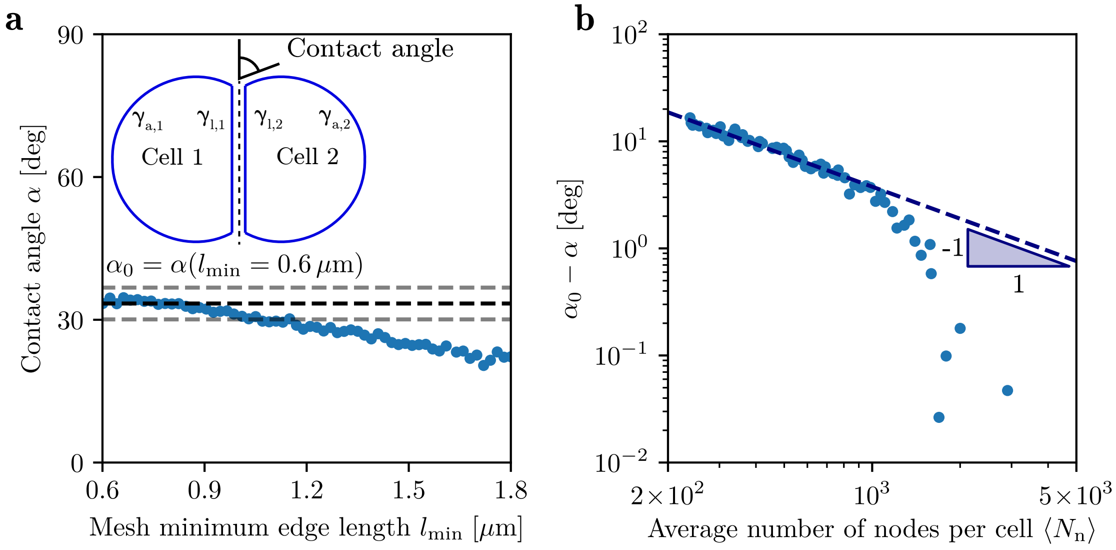


**Figure S4: Validation of the spring-based contact model in mechanical equilibrium. A**, Contact angle $\alpha$ between a pair of cells as a function of the minimum mesh edge length $l_{min}$. $l_{min} = 0.6\,\mu m$ corresponds to a high mesh resolution with $\approx 3,500$ nodes per cell, while $l_{min} = 1.8\,\mu m$ corresponds to a low mesh resolution with $\approx 350$ nodes per cell. The apical surface tensions of the cells ($\gamma_{a,1}$ and $\gamma_{a,2}$) as well as their lateral surface tensions ($\gamma_{l,1}$ and $\gamma_{l,2}$) were set to $2.5 \times 10^{-4}$ N/m in all simulations. The adhesion strength between the cells was kept constant at $\omega = 2.5 \times 10^{-4}$ Pa/m in all simulations. The black dotted line represents the contact angle value $\alpha_0$ obtained in a simulation with high mesh resolution ($l_{min} = 0.6\,\mu m$), while the grey dotted lines correspond to a $\pm 10\%$ variation of this value. **B**, Difference in contact angle with respect to a high resolution simulation as a function of the average number of nodes per cell. The dark blue dashed line shows the relationship $\alpha = \alpha_0 - a/\langle N_n \rangle$ with fitted coefficient $a = 3.6 \times 10^3$ deg.

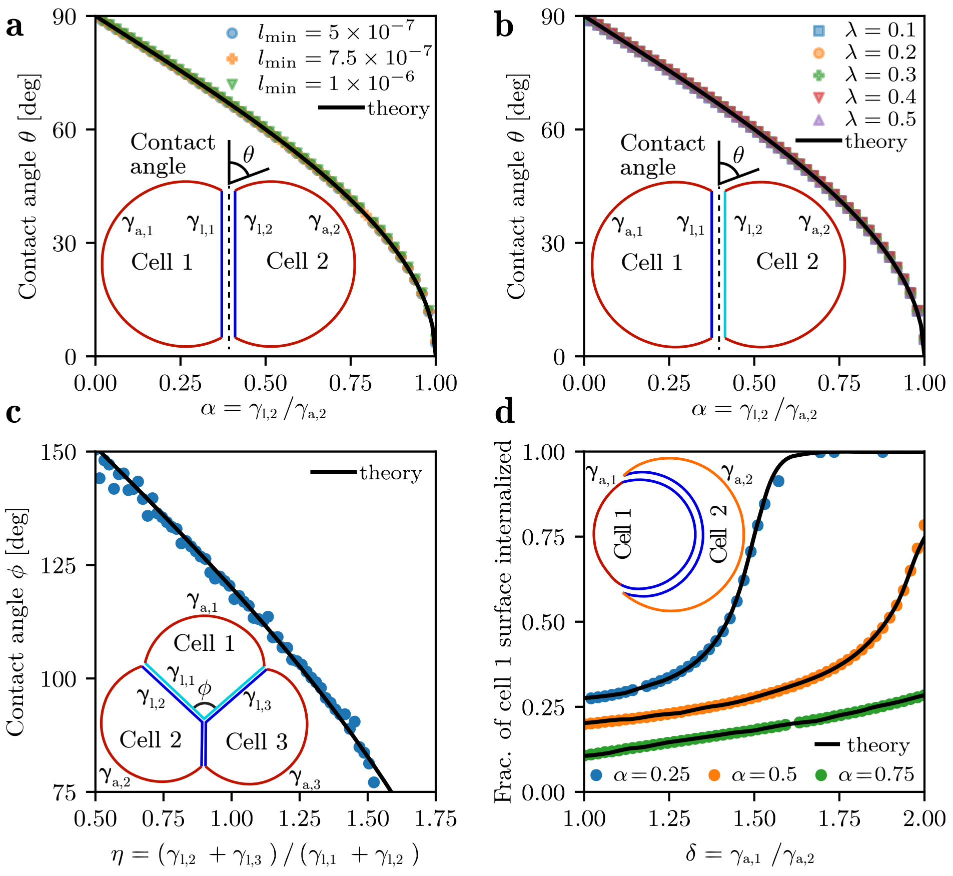


**Figure S5: Validation of the coupling-based contact model in mechanical equilibrium.** **A**, Contact angle $\theta$ between a pair of cells as a function of their lateral ($\gamma_{l,i}$) to apical ($\gamma_{a,i}$) surface tension ratio, and mesh resolution ($l_{\min}$). $\gamma_{a,1} = \gamma_{a,2}$, and $\gamma_{l,1} = \gamma_{l,2}$. The theoretical black curve is given by the Young–Dupré equation as $\cos\theta = \gamma_{l,2}/\gamma_{a,2}$. **B**, Contact angle $\theta$ between a pair of cells as a function of their lateral to apical surface tension ratio. $\gamma_{a,1} = \gamma_{a,2}$, but $\gamma_{l,1} \neq \gamma_{l,2}$. $\lambda$ is the deviation factor of each cell lateral surface tension from the mean lateral surface tension *i.e.* $\gamma_{l,1} = \overline{\gamma}(1-\lambda)$, and $\gamma_{l,2} = \overline{\gamma}(1+\lambda)$ where $\overline{\gamma} = (\gamma_{l,1} + \gamma_{l,2})/2$. The theoretical black curve is given by $\cos\theta = \overline{\gamma}/\gamma_{a,2}$. **C**, Contact angle $\phi$ at a tricellular junction as a function of the ratio $\eta = (\gamma_{l,2} + \gamma_{l,3})/(\gamma_{l,1} + \gamma_{l,2})$. $\gamma_{a,1} = \gamma_{a,2} = \gamma_{a,3}$, and $\gamma_{l,1} \neq \gamma_{l,2} = \gamma_{l,3}$. The theoretical line is obtained from the Young–Dupré law and follows $\cos(\phi/2) = \eta/2$. **D**, Proportion of cell 1 surface area internalized as a function of the apical surface tension ratio $\delta = \gamma_{a,1}/\gamma_{a,2}$. $\gamma_{l,1} = \gamma_{l,2}$, and $\alpha = \gamma_{l,2}/\gamma_{a,2}$. The theoretical curves were calculated based on the Lagrangian approach presented in [1].

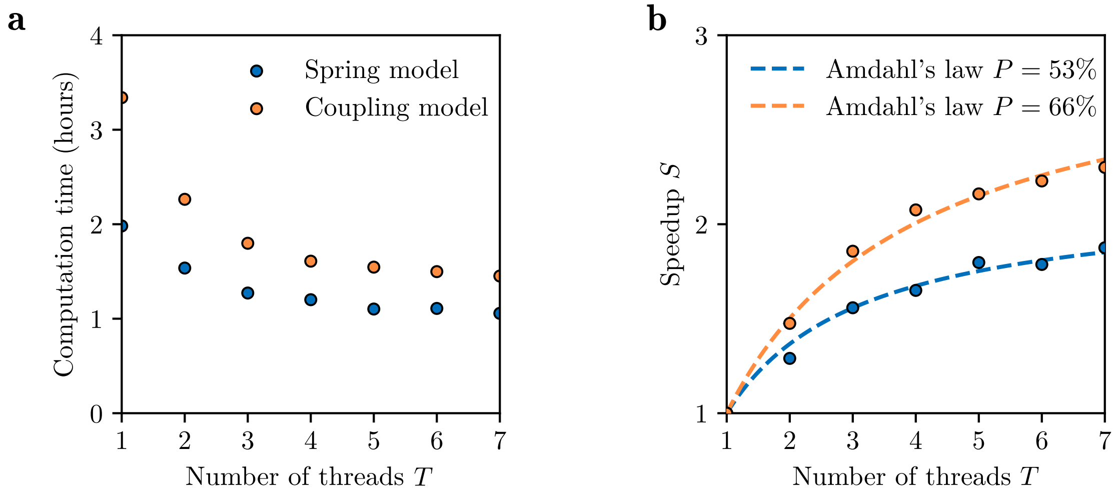


**Figure S6: Parallel computational performance.** **A**, Computation time required to simulate the development of a tissue from 1 to 500 cells with respect to the number of threads ($T$). The computation time was recorded for both contact models available in SimuCell3D. The average number of nodes per cell in these simulations was approximately 700. **B** Speedup $S$ as a function of the number of threads. Amdahl's law: $S = 1/(1 - P + P/T)$, where $P$ is the parallel fraction. All simulations were performed on an Intel Xeon W-2125 processor (8 cores, 4.5 GHz).
